# Structure based virtual screening identifies novel competitive inhibitors for the sialoglycan binding protein Hsa

**DOI:** 10.1101/2020.03.27.006247

**Authors:** Rupesh Agarwal, Barbara A. Bensing, Dehui Mi, Paige N. Vinson, Jerome Baudry, Tina M. Iverson, Jeremy C. Smith

## Abstract

Infective endocarditis (IE) is a cardiovascular disease often caused by bacteria of the *viridans* group of streptococci, which includes *Streptococcus gordonii* and *Streptococcus sanguinis*. Previous research has found that a serine-rich repeat (SRR) proteins on the *S. gordonii* bacterial surface play a critical role in pathogenesis by facilitating bacterial attachment to sialyated glycans displayed on human platelets. Despite its important role in disease progression, there are currently no anti-adhesive drugs available on the market. Here, we performed structure-based virtual screening using an ensemble docking approach followed by consensus scoring to identify novel inhibitors against the sialoglycan binding domain of the SRR adhesin protein Hsa from the *S. gordonii* strain DL1. *In silico* cross screening against the glycan binding domains of closely related SRR proteins from five other *S. gordonii* or *S. sanguinis* strains was also performed to further reduce false positives. Using our *in silico* screening strategy we successfully predicted nine compounds which were able to displace the native ligand (sialyl-T antigen) in an *in vitro* assay and bind competitively to adhesin protein Hsa (∼20% hit rate).

## Introduction

Infective endocarditis (IE) (or bacterial endocarditis (BE)) is a life-threatening cardiovascular infection of the inner lining of the heart muscle (endocardium) and heart valves^1^. The *viridans* group of streptococci account for ∼17-45% of all cases of IE^2, 3^. If the infection remains untreated, these bacteria form a biofilm that eventually destroys the valves and result in heart failure^4^. Moreover, these bacteria may also form small clots (emboli) which can block small arteries. IE affects 10,000-20,000 patients in the US every year and is associated with an in-hospital mortality rate of ∼20% and a five year mortality rate of ∼ 40-70 %^1^. Currently, the most common treatment for endocarditis combines surgical intervention with antibiotics. The rise in antibiotic resistance^5^ has limited our pharmacological options^6, 7^, and resistant organisms have increased the mortality rate^8^. Although surgical intervention physically removes the biofilm from the heart valves, 47% or more of the patients eventually require valve replacement due to the damage incurred^9^. Given the associated morbidity and rising mortality rate, there is an urgent need to develop novel therapies against IE.

Previous studies have reported that the binding of bacteria to host platelets contributes to the colonization of damaged aortic valves^4^. A cell wall-anchored serine-rich repeat (SRR) protein mediates the adherence of *S. gordonii* and *S. sanguinis* to sialoglycans displayed on the human platelet^10^ glycoprotein GPIb^11, 12^. SRR proteins have been demonstrated to be virulence factors for endocarditis^11, 12^, and disrupting the interaction between the SRR protein and sialoglycans on host platelets may therefore reduce virulence. *Streptococcus gordonii* is one of the well-studied species that cause IE and is a normal component of the human oral microbiota^13^. Platelet binding by *S. gordonii* strains M99 and DL1 are facilitated by the homologous SRR proteins GspB and Hsa, respectively^14^. Although these two share significant sequence identity, their ligand binding regions (BRs) differ significantly and have different sialoglycan selectivity^11, 15, 16^. GspB binds with narrow selectivity to sialyl-T antigen (sTa) whereas Hsa binds promiscuously to a range of glycans^11, 15, 16^. Anti-adhesive therapies have been explored for the treatment of a wide range of other bacterial infections^17^, but have not yet been pursued for IE. Anti-adhesives can, in principle, complement traditional antibiotics and improve their efficacy, potentially eliminating the need for surgical intervention.

The crystal structures of the BRs from a number of *S. gordonii* and *S. sanguinis* SRR proteins have been solved^18-20^. These all have two domains which are associated with sialoglycan binding: the Siglec (Sialic acid-binding immunoglobulin-like lectin) domain and the Unique domain (for which function is not known completely). Furthermore, recent studies have identified that the three loops (CD, EF and FG) adjacent to the sialoglycan binding site are critical for the affinity and selectivity between ligands^20^. Additionally, it has been reported that a conserved “YTRY” motif in the binding site is necessary for formation of hydrogen bond interactions with the sialic acid of the native ligand^20^. Importantly, there are also human sialoglycan-binding proteins, that contain a sialoglycan binding site but are distinct from the bacterial sialoglycan binding adhesin proteins^21^.

With the above structural and dynamic information, a structure-based drug design (SBDD) strategy was followed to target SRR adhesin proteins. In our SBDD pipeline, we targeted the BR of the well-characterized SRR protein Hsa (Hsa_BR_), using *in-silico* virtual screening. Moreover, since Hsa_BR_ binds promiscuously to many glycans using a conformation selection mechanism^20^, it is potentially a good target for SBDD. Here, instead of using only the crystal structure for SBDD, we used molecular dynamics (MD) simulation to capture the flexibility of the binding pocket and generate an ensemble of protein conformations^22^. Following subsequent high throughput ensemble docking, we prioritized the compounds using consensus scoring, which has previously shown to reduce the number of false positives and increase the hit rate^23^. To further improve our predictions, we cross screened the compounds against the BRs from five Hsa homologues, and identified compounds which bound to Hsa_BR_ with relatively higher docking scores compared to other BRs. From our virtual screening predictions, we were able to achieve a high hit rate of ∼20%, finding that 9 out of 50 compounds that were suggested for experimental validation were able to displace the native ligand from the Hsa_BR_ binding pocket. To our knowledge, these are the first pharmacological compounds known to inhibit binding by the SRR adhesin protein.

## Methods

### System preparation and molecular dynamics simulation

Crystal structures of the sialoglycan binding proteins Hsa_BR_ (PDB 6EFC)^20^, GspB_BR_ (PDB 6EFA)^20^, 10712_BR_ (PDB 6EFF)^20^, SK150_BR_ (PDB 6EFB)^20^, SrpA_BR_ (PDB 5EQ2)^18^, and SK678_BR_ (PDB 6EFI)^20^ were used in this study. Molecular dynamics (MD) simulations was performed on all these proteins using the Amber14 ff14SB force-field parameters^24, 25^. Each of these proteins was surrounded by an octahedral box of water model TIP3P^26^ of 10 Å. First, the protein structure was held fixed with a force constant of 500 kcal mol^-1^ Å^-2^ while the system was minimized with 500 steps of steepest descent followed by 500 steps with the conjugate gradient method. In the second minimization step, the restraints on the protein were removed and 1000 steps of steepest descent minimization were performed followed by 1500 steps of conjugate gradient. The system was heated to 300 K while holding the protein fixed with a force constant of 10 kcal mol^-1^ Å^-2^ for 1000 steps. Then, the restraints were removed, and 1000 MD steps were performed. The SHAKE algorithm^27^ was used to constrain all bonds involving hydrogen in the simulations. 200 ns production MD was performed at 300 K using the NPT ensemble and a 2 fs time step with nonbonded cutoff of 10 Å. The temperature was fixed with the Langevin dynamics thermostat^28^ and the pressure was fixed with a Monte Carlo barostat^29^. This procedure yielded a total of 20,000 snapshots for subsequent analyses. Three independent runs were performed for each simulation.

### In silico screening

Ensemble docking is an *in-silico* structure-based drug discovery method using an ‘ensemble’ of drug target conformations to discover novel inhibitors^22^. The workflow used is shown in Fig. 1. The ensemble was constructed by clustering snapshots from molecular dynamics (MD) simulation trajectories by root mean square deviation (RMSD) of the binding pocket residues and loops (**Table S1**) surrounding the binding pocket with the hierarchical agglomerate clustering algorithm using Cpptraj module^30^.

**Figure 1:**
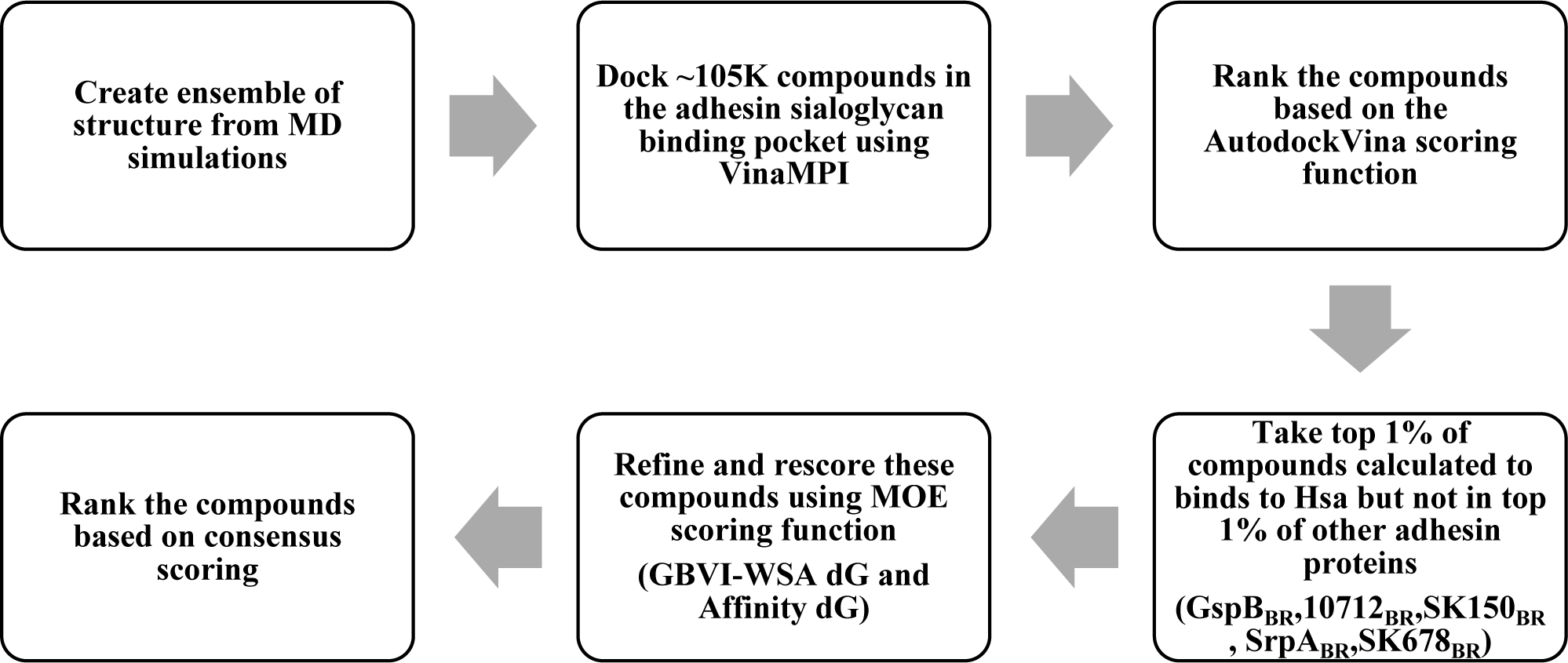
Structure based virtual screening strategy workflow.

The Vanderbilt small molecule database (∼105,000 compounds) was docked to an ensemble of 5 conformations (4 representative structures obtained from clustering from MD and 1 crystal structure) with a cubic box with edges of ∼30 Å. VinaMPI^31^, a parallel version of AutodockVina^32^, was used to perform the *in silico* screening. The docked poses were then ranked by the AutodockVina scoring function^33^. The compounds were not only screened for Hsa_BR_ but also cross screened with 5 adhesin proteins (GspB_BR_, 10712_BR_, SK150_BR_, SrpA_BR_, SK678_BR_). In this study, we aim to find a potential new scaffold and since all the adhesin proteins natively bind to a common motif (sialic acid), we believe the compounds which bind to all the adhesin proteins (with a high score) are more likely mimicking sialic acid or are very promiscuous. Hence, cross screening was performed to remove these sialic acid mimetics and promiscuous compounds, thus reducing the number of false positives.

From this ranked list of compounds, we tested compounds which were within the top 1% for Hsa_BR_ but not within the top 1% of the other 5 BRs (GspB_BR_, 10712_BR_, SK150_BR_, SrpA_BR_, SK678_BR_).

We note that we only experimentally tested binding to Hsa _BR_ and not the selectivity of the predicted binders. Next, the resulting ∼250 compounds were refined and rescored using two MOE scoring functions^34^. The non-polar hydrogens (not included in Vina docking protocol) were added before performing the “induced fit” docking protocol in MOE^34^. The docking poses were ranked using GBVI-WSA dG and Affinity DG scoring functions^34^. Using consensus scoring, the top 50 compounds, which were in top 1% in all the scoring functions, were suggested for experimental validation. A flowchart of the screening methodology used is shown in **Fig. 1**.

### Cheminformatics

All the physicochemical properties and fingerprints of small molecules were calculated using combination of MOE^34^, ChemBioServer^35^ and RDkit^36^. MACCS fingerprints were calculated for each compound and similarity between them were compared with the Tanimoto coefficient, followed by hierarchical clustering to cluster the similarity matrix.

### Experimental assays

#### Protein expression and purification

GST-tagged Hsa_BR_ was expressed and purified as described in ref^16^. GST-Hsa_BR_ was expressed under the control of the pGEX-3X vector in *E. coli* BL21 (*DE3*) in a Terrific Broth medium with 50 µg/ml kanamycin at 37 °C. When the OD_600_ reached 1.0, expression was induced with 1 mM IPTG at 24 °C for 5 hrs. Cells were harvested by centrifugation at 5,000 × *g* for 15 min, optionally washed with 0.1 M Tris-HCl, pH 7.5, and stored at –20 °C before purification. The frozen cells were resuspended in homogenization buffer (20-50 mM Tris-HCl, pH 7.5, 150-200 mM NaCl, 1mM EDTA, 1 mM PMSF, 2 µg/ml Leupeptin, 2 µg/ml Pepstatin) then disrupted by sonication. The lysate was clarified by centrifugation at 38500 × *g* for 35-60 min and passed through a 0.45 µm filter. Benchtop purification was performed at 4 °C using Glutathione Sepharose 4B beads, with pure GST-Hsa were eluted with 30 mM GSH in 50 mM Tris-HCl, pH 8.0.

#### AlphaScreen high-throughput screening assay

We used the AlphaScreen modification of an ELISA as the primary target-based proximity assay to monitor ligand displacement. AlphaPlate (Cat # PE 6005351, Lot # 8220-16081) with 384-well was used for the screening. In the experimental setup, biotinylated sialyl T antigen (sTa) was coupled to a streptavidin donor bead and GST-tagged Hsa was coupled to an anti-GST conjugated acceptor bead in PBS (phosphate buffered saline). The reaction was excited at 680 nm to stimulate singlet oxygen-mediated energy transfer to the acceptor bead, which can be detected at 615 nm. The dose-dependent signal reflects the number of bead-coupled adhesins bound to bead-coupled glycans.

The GST-Hsa_BR_ concentration was titrated in a 10 point-3 fold dilution starting from 1000 nM. The biotinylated sTa concentration was titrated in a 9 point-3 fold dilution starting from 100 nM, The hooking point was found to be 3 nM for GST-Hsa_BR_ and 3 nM for biotinylated sTa. The final chosen concentrations used in the screen was 2 nM of biotinylated sTa and 1 nM of GST-Hsa_BR_.

The untagged-sTa competition assay was run in a dose response mode. Untagged-sTa was tested in an 11 point-3 fold CRC, starting from 30 μM as the (1x) concentration. 30 μM was finally used in the screen.

We applied this assay to the evaluation of the test compounds that were predicted as binding to Hsa_BR_ using virtual screening. This initial screen was performed with all test compounds in duplicate at a final concentration of 10 μM and DMSO was used as the negative control and unbiotinylated sTa was used as a positive control.

#### Z’ factor calculation

The Z’ factor is an indicator of high throughput screening assay performance and was calculated as follows:

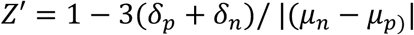

The standard deviations and means of the positive (p) and negative (n) controls are denoted by *δ*_*p*_, *μ*_*p*_ and *δ*_*n*_, *μ*_*n*_ respectively. DMSO and untagged sTa are the positive and negative control respectively.

#### Hit identification analyses

The alpha value of each test compound was measured and was filtered using 1-fold, 2-fold or 3-fold of either standard deviation (SD) from the mean of the entire test compound group, or absolute deviation from the median (MAD) of the entire population. Compounds that satisfied any of these criteria were considered for the next round of filtering using percentage control (PC), calculated as follows:

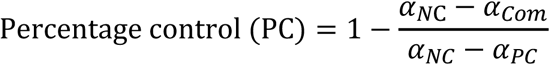

where, *α* is the average alpha value for negative control (*α*_*NC*_), positive control (*α*_*PC*_) and compounds tested (*α*_*Com*_). PC is a measure of the alpha signal of the 10 μM test compound in percentage of the controls.

## Results and Discussion

### I. Protein dynamics and conformations

We used MD simulations to capture the internal dynamics of the proteins and find binding site conformations not seen in the crystal structure^37^. We calculated the root mean square fluctuation (RMSF) to identify the flexible regions (**Fig 2a**). Although the overall structure of the Siglec domain is rigid, we observed that the loops (CD, EF and FG) close to the binding pocket are flexible for all the adhesin proteins (**Fig 2a, S1)**. In the case of Hsa_BR_, we observed that the CD and EF loops constitute the most flexible region of the protein. Moreover, critical binding pocket residues other than in these loops were identified from the crystal structure of Hsa_BR_ and the native ligand (sTa) (**Table S1**).

**Figure 2:**
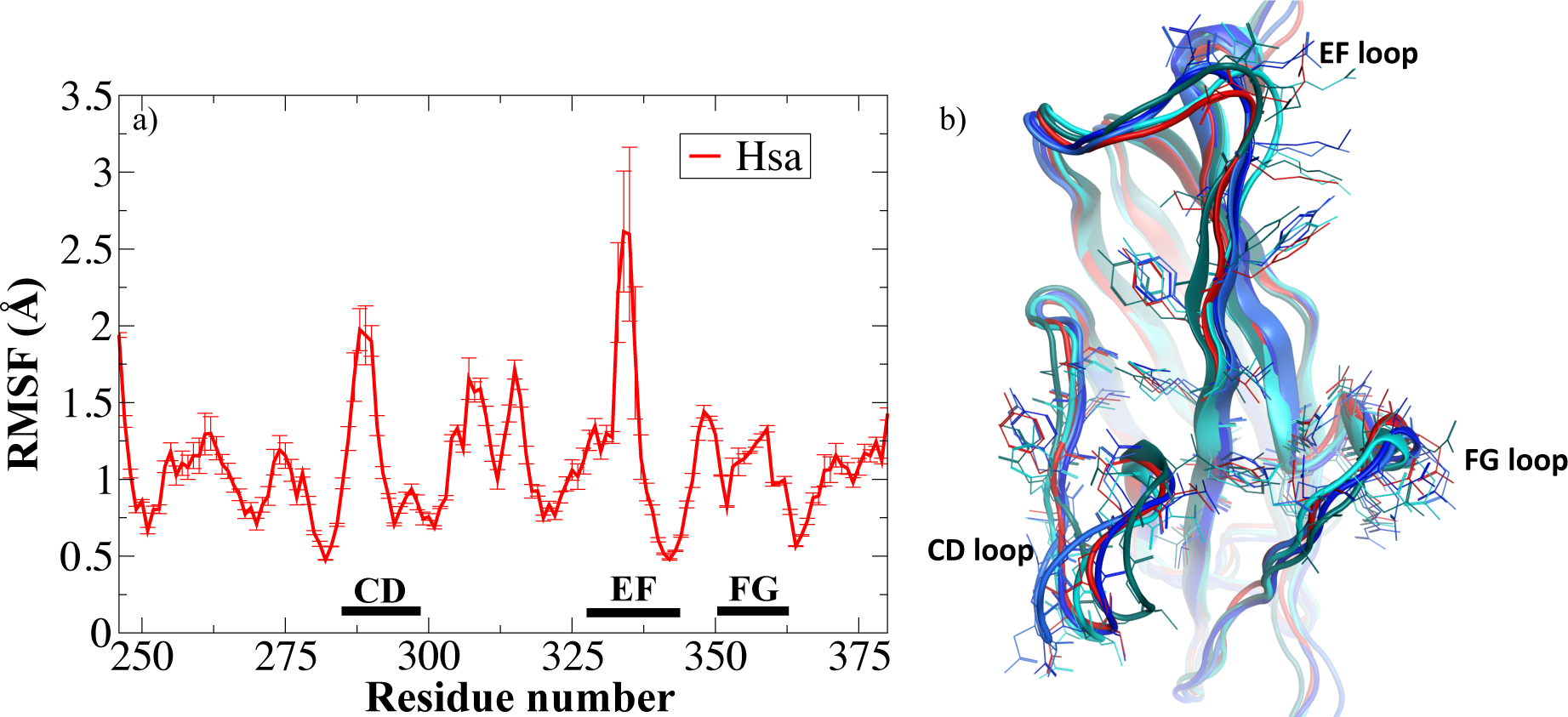
a) Root mean square fluctuation of Hsa_BR_ from MD simulation showing DC, EF, FG loop regions; b) Superimposed structures (in ribbon) of Hsa_BR_ obtained from different clusters showing residues (in stick) used to during the clustering: crystal structure (in red) and ensemble structure (in shade of blue).

To capture new conformations of the binding pocket that deviate from the initial crystal structure, the root mean square deviation (RMSD) of the loop residues and other critical residues (previously known to bind to the native ligand) (**Table S1**) were used to cluster the MD trajectories. The clustering resulted in four different clusters. The structure closest to the centroid of each cluster was used for docking. The “ensemble” of structures obtained from clustering and crystal structure were superimposed to observe the deviation of structures (**as shown in Fig 2b)**. We observe that all the structures had similar secondary and backbone structures and the RMSD_Calpha_ of the Siglec domain was calculated to be within ∼1.5 Å. However, as seen in the superimposed structures (**Fig 2b**), the loop regions (especially CD and EF loop) have different orientations in the “ensemble” when compared to the crystal structure. Similarly, we observed that the side chains in the binding pocket residues orient differently between the structures, which can be critical for rigid body docking.

### II. Physicochemical properties of small molecule database

The five structures obtained from MD simulations and the existing crystal structure were screened against the Vanderbilt small molecule database containing ∼102K compounds. However, before performing the virtual screening, we wanted to characterize the physicochemical diversity of the small molecule database. Firstly, we calculated the molecular weight (MW) of the compounds (**Fig 3a**), which is known to be critical for safety and tolerability reasons^38^. The Vanderbilt database has compounds with MW less than 500 Da that are considered to improve druglikeness^39, 40^ and also has low MW compounds (<300 Da) that are considered better initial hits because they serve as effective chemical starting points for lead optimization^41^. The polar surface area and the number of rotatable bonds have been found to better discriminate between compounds that are orally active. It has been predicted that compounds with 10 or fewer rotatable bonds and those having a polar surface area of less than 140 Å^2^ have a good oral bioavailability^42^. In our database, we observed that most compounds had a mean polar surface area of ∼150 Å^2^ and less than 10 rotatable bonds (**Figs 3b, c**). Lipophilicity (SLogP) is another factor which is known to influence drug potency, pharmacokinetics, and toxicity^39, 43^. Compounds with SLog P values between −0.4 to +5.6 range are known to be more “druglike”^40, 44^. Here, we found that most of the compounds fall within this range (**Fig 3d**). Although the above is a set of physicochemical properties that are considered to be important for different aspects of druggability, there have been numerous FDA approved drugs which violate one or more of these rules^45, 46^.

**Figure 3:**
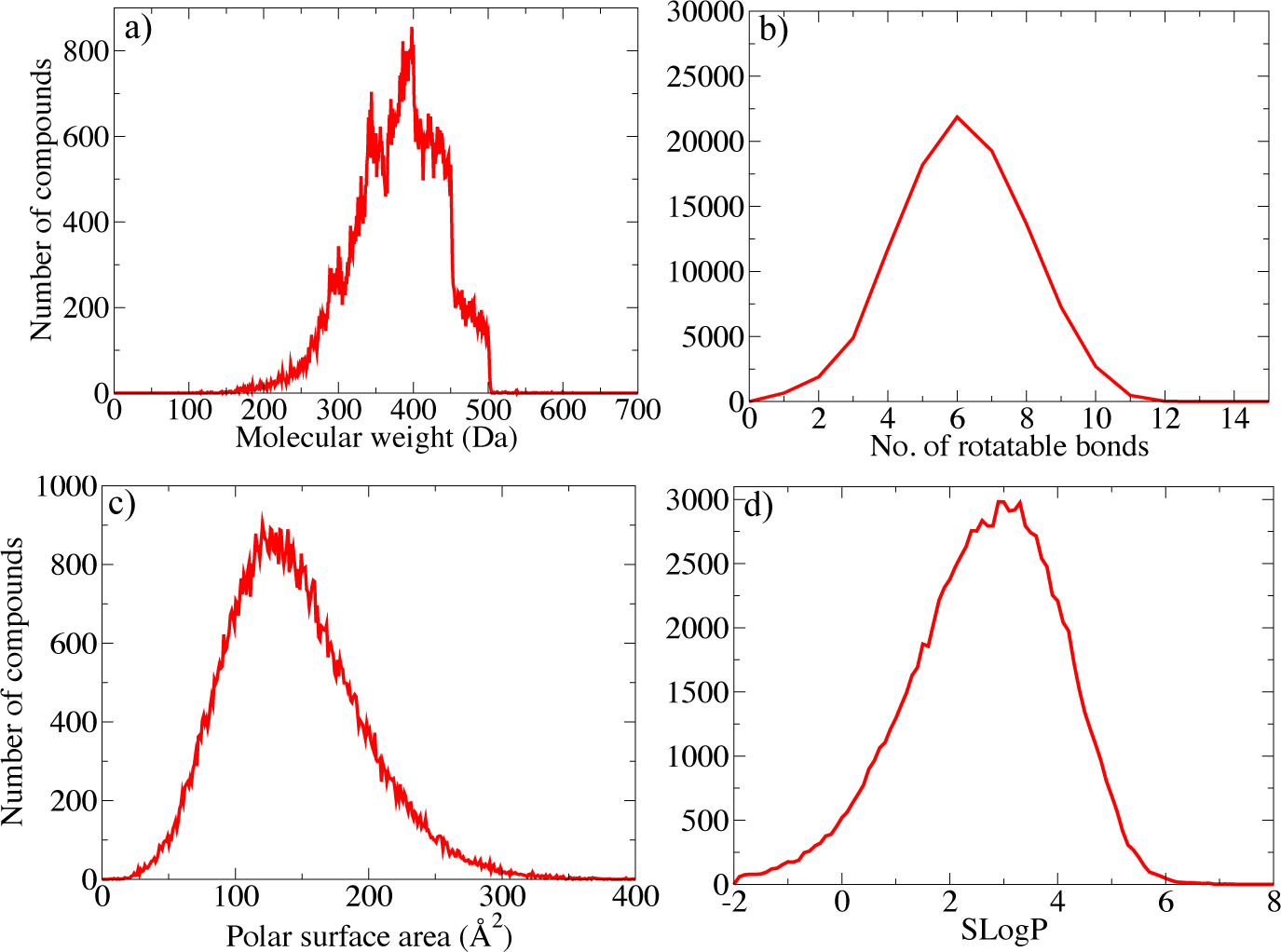
Density profile of physicochemical properties of small molecule database: molecular weight (a), number of rotatable bonds (b), polar surface area (c), and Log of the octanol/water partition coefficient: SLogP (d).

### III. Docking results and poses

After our virtual screening, we first ranked all the top poses for each compound based on the Autodock Vina scoring function^33^. Subsequently, we selected those compounds that were in the top 1% for Hsa_BR_ but did not rank within the top 1% for any other adhesin protein (GspB_BR_, 10712_BR_, SK150_BR_, SrpA_BR_, SK678_BR_). This was followed by implementing consensus scoring in which the poses (obtained from AutodockVina) were energy-minimized and then rescored using two MOE scoring functions^34^ (as mentioned in the Methods section). In the end, compounds that ranked within the top 50 for all the three scoring functions were suggested for experimental validation. Since the goal of this work was to find competitive inhibitors, we calculated the number of compounds that formed at least one hydrogen bond (HB) with one of the residues known to bind to the native ligand. Backbone atoms of residues Tyr 337 and Thr 339 of Hsa_BR_ are known to form HBs with the native ligand (sTa) (**Fig. 4b**), whereas sidechains of residues Arg 340, Val 285 form multiple HBs with sTa. All the top 50 compounds, from our strategy were found to form at least one HB with the residues known to bind sTa (**Fig. 4a**). Moreover, out of these 50 compounds, 25 compounds (50%) were predicted to bind to one of the “ensemble” structures generated from MD simulations and not to the crystal structure. This further illustrates the usefulness of using ensemble docking.

**Figure 4:**
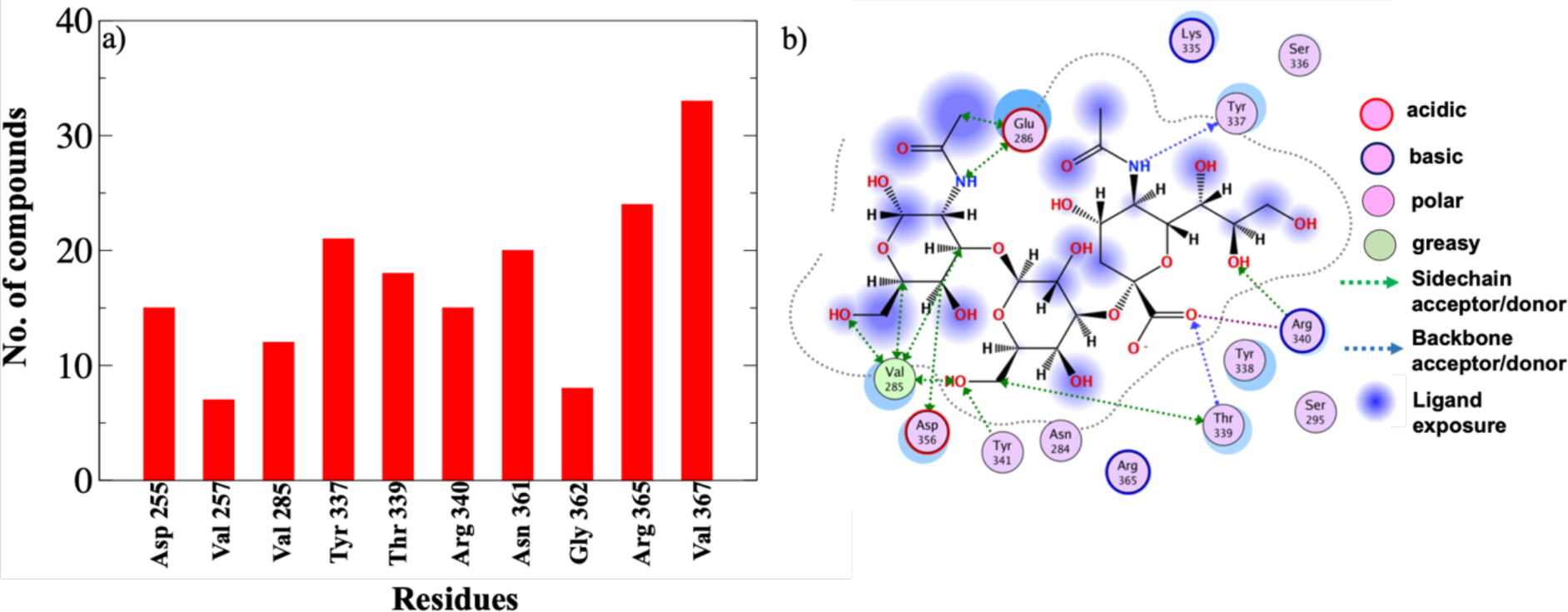
a) Number of compounds within the top 50 compounds interacting with key residues in the Hsa_BR_ binding pocket; b) interaction map of native ligand (sTa) from crystal structure.

### IV. Experimental validation

Alpha assay screening was performed for the top 50 compounds predicted to displace sTa (the highest affinity native ligand) from Hsa_BR_. The Z’ factor value of the DMSO (negative control) versus untagged sTa (positive control) was 0.32. After filtering the small molecules using the experimental data based on the percentage control (PC), nine hits were retained. These nine compounds showed a statistically significant decrease in the signal when the two replicates were averaged (**Fig. 5**). These compounds have a PC three standard deviations outside the mean of the negative control (DMSO).

**Figure 5:**
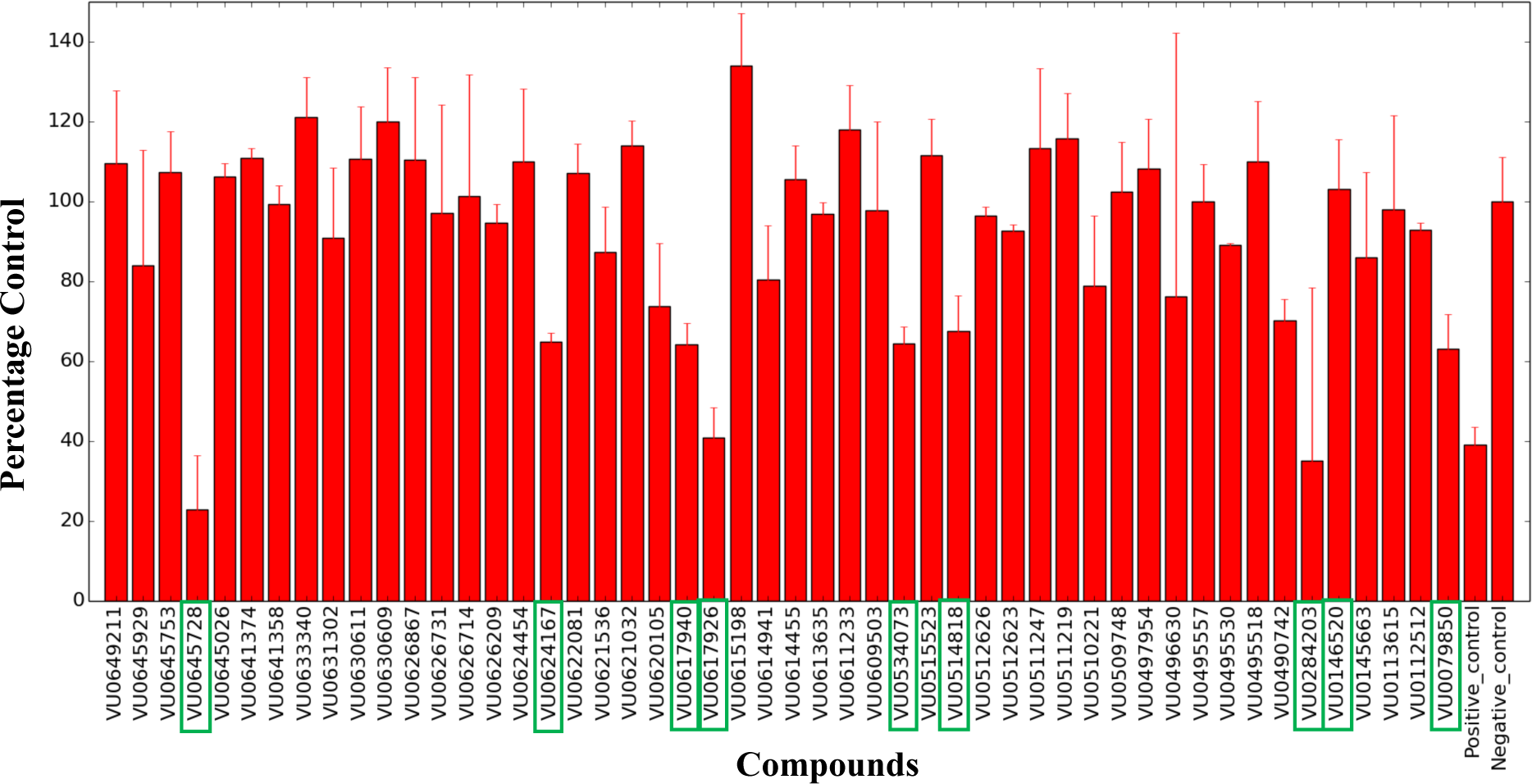
Alpha Screen assay. Hits are marked by green boxes on the X-axes. Error bar represent the standard deviation.

The IC_50_ of the untagged sTa (positive control) was calculated as 8.67 μM (**Fig. S2a**). At 10μM concentration, the PC of the untagged sTa was 39% (**Fig. S2b**). At the same concentration, the PC of the 9 hits ranges from 23% to 70% and, out of these, two compounds have PC values of less than 39% and one has a PC of 41% (**Table S2)**.

### V. Computational and binding pose analyses of validated hits

The nine hit compounds were screened for 25 known toxic and carcinogenic fragments, such as anthracene, quinone, hydroquinone, butenone--Michael acceptor, chloroethane--Michael acceptor^35^. Of the 9 experimentally validated compounds (C1-C9) (**Table 1**), only Compound 1 (C1) was identified as potentially toxic, containing a benzo-dioxane and a catechol group. Moreover, to test the similarity between these hits and the native ligand, fingerprint-based hierarchical clustering was performed. We found four clusters (**as shown in Fig. S3**), which showed that the compounds identified from the screen are diverse among themselves and are not similar to the native ligand. Additionally, we also tested the compounds for Lipinski’s rule^40^, to evaluate druglike-ness of the compounds. C4 was the only molecule with one violation (with 11 hydrogen bond acceptors), whereas all the other compounds satisfied all the 4 rules.

**Table 1:**
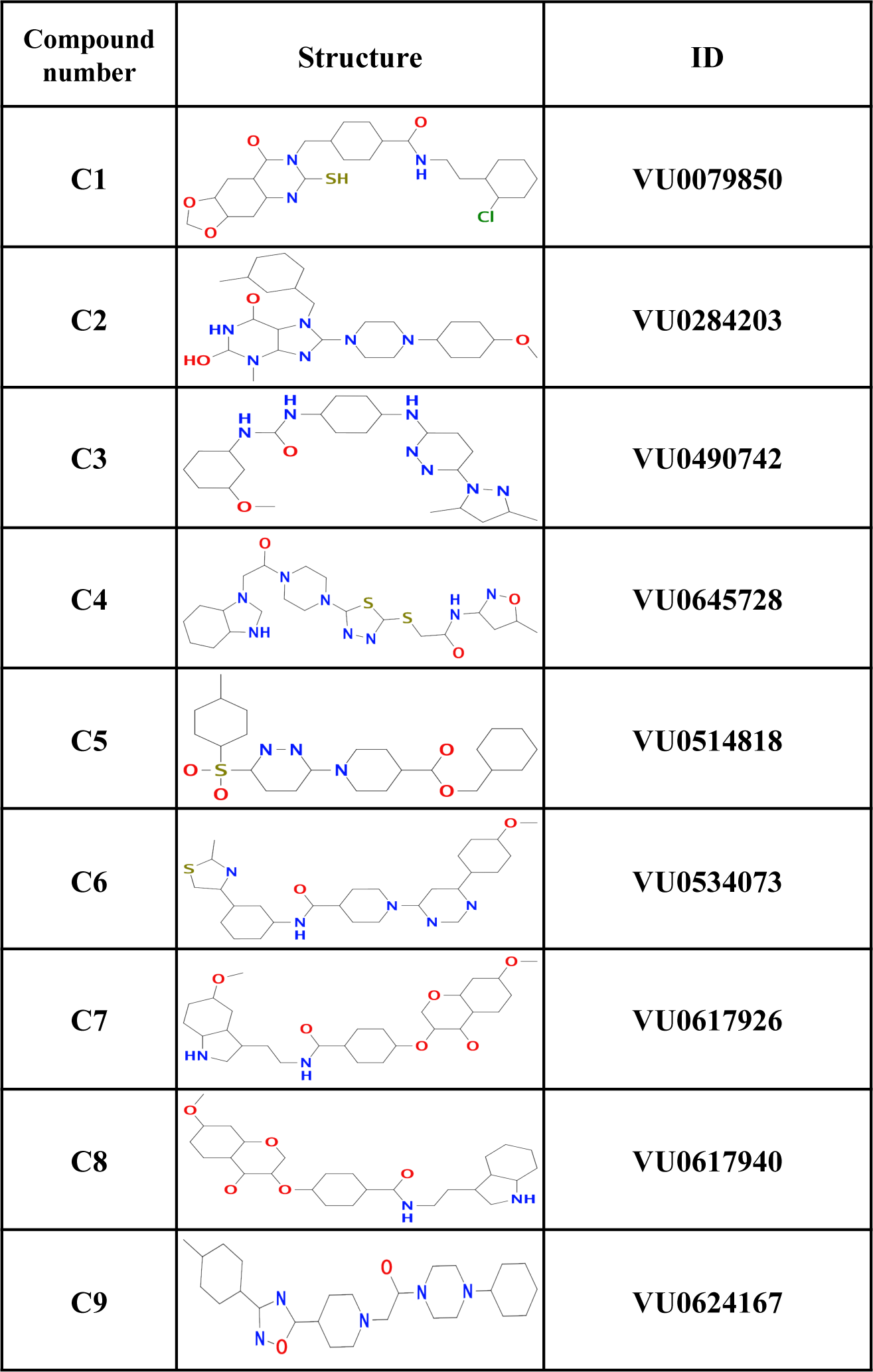
The compound number, structure and ID of the nine validated hits.

| Compound number | Structure | ID |
| --- | --- | --- |
| C1 |  | VU0079850 |
| C2 |  | VU0284203 |
| C3 |  | VU0490742 |
| C4 |  | VU0645728 |
| C5 |  | VU0514818 |
| C6 |  | VU0534073 |
| C7 |  | VU0617926 |
| C8 |  | VU0617940 |
| C9 |  | VU0624167 |

Following the above cheminformatics analyses of the experimentally-validated hits, we examined the computational models of the best poses and the interactions of the nine hits shown to have inhibition experimentally (**Figs. 6, S4 and table S3**). In the models C1, C2, C4, and C5 form backbone HBs with Asp 255 and compounds C2-4 form backbone HBs with Val 367 while other compounds (C1, C5-9) form side chain HBs with Val 367. Other residues that form HBs with most of the compounds are Thr 339, Val 285 and Asn 361 (**Table S3)**. All these residues form HBs with the native ligand or are in close proximity of the native ligand. Hence, it is likely that these nine compounds are able to displace the native ligand in part because they form HBs with these critical residues.

**Figure 6:**
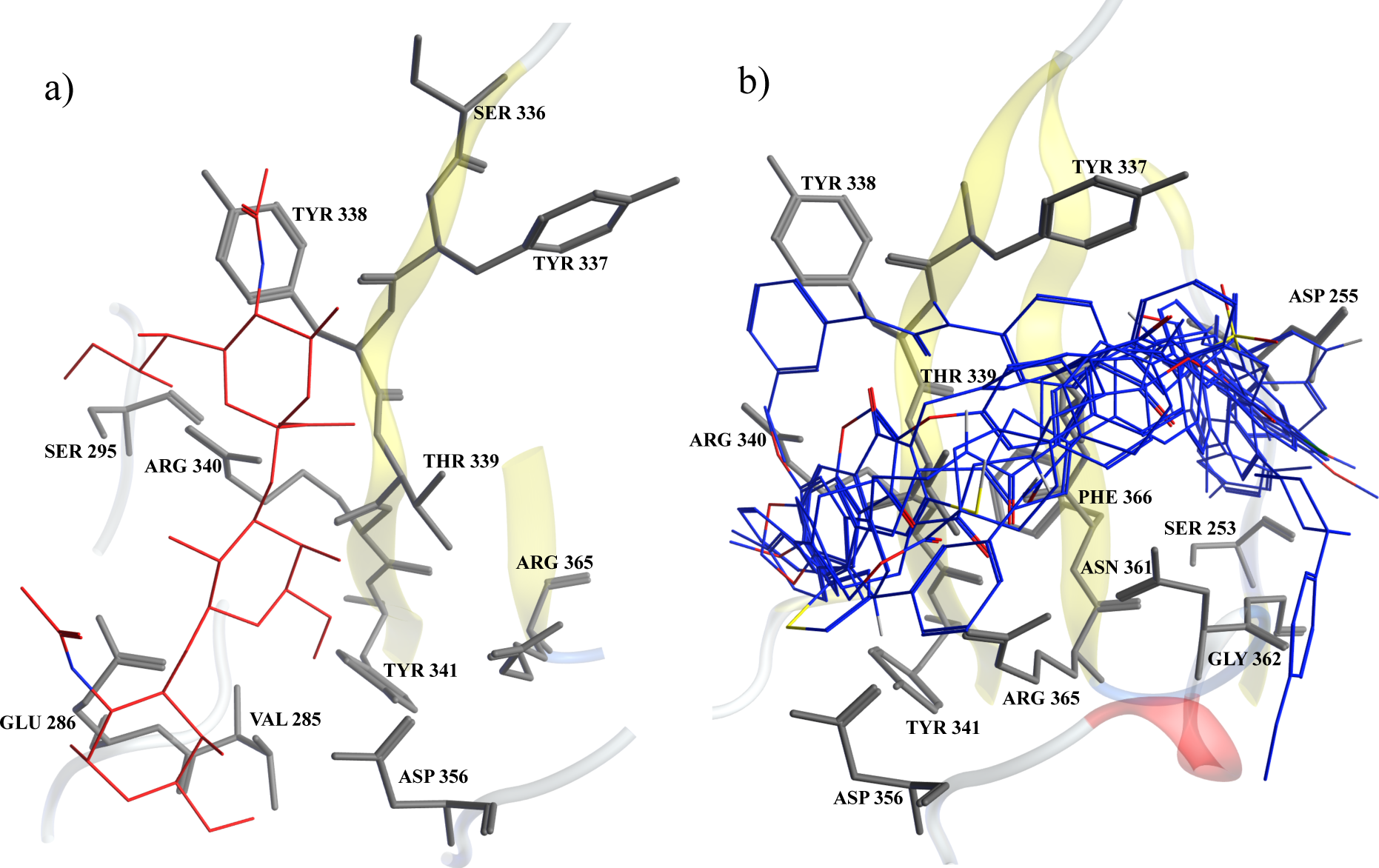
Docking pose of a) native ligand (sTa) (in red) and b) 9 validated compounds (in blue) in the binding pocket of Hsa_BR_ showing residues (in grey).

## Conclusions

The SRR protein Hsa has been considered an attractive molecular target for drug development due to its role in infective endocarditis (IE). It is noteworthy that there is no vaccine or anti-adhesive drug approved against IE. Here, we performed structure-based virtual screening to identify competitive inhibitors for Hsa_BR_. We combined three different SBDD strategies; ensemble docking, cross screening, and consensus scoring in one pipeline. For the ensemble docking, we generated an ensemble of receptor conformations from MD simulation, and then cross screened against five homologs (GspB_BR_, 10712_BR_, SK150_BR_, SrpA_BR_, SK678_BR_). In the last step, three scoring functions (AutodockVina^33^ and MOE^34^) were used to rank and prioritize the list of compounds. The Vanderbilt database was used for the small molecules since it covers a wide distribution of different physicochemical properties.

The goal of combining these strategies was to improve the hit rate and reduce the number of false positives. Indeed, we were able to achieve a hit rate of ∼20% and identified nine compounds that could displace the native ligand in the experimental assay. The binding poses of all the nine compounds identified from docking show that they are in close proximity with residues known to form HBs with the native ligand (sialyl-T antigen). These compounds may be used as a starting point for further medicinal chemistry optimization. Further studies need to be conducted to characterize the binding affinity and pose of these identified compounds, and similar analyses for other sialoglycan-binding SRR proteins is ongoing.

## Acknowledgements

We would like to thank Dr. Jarrod Smith for providing the small molecule database. This work was supported in part by National Institutes of Health (NIH) grant AI106987 (TMI).

## Notes

#### Summary of Updates

The author's (Paige) name was wrongly spelled.

