## Supplementary information for "Structure based virtual screening identifies novel competitive inhibitors for the sialoglycan binding protein Hsa"

### Table S1: Residues included during clustering of MD simulation snapshots to form an 'ensemble'

| Protein | Residues |
|---|---|
| **$Hsa_{BR}$** | 285-295, 333-343, 356-365 |
| **$SK678_{BR}$** | 297-307, 342-352, 366-376 |
| **$SrpA_{BR}$** | 288-298, 338-348, 363-368 |
| **$10712_{BR}$** | 286-299, 335-345, 355-366 |
| **$GspB_{BR}$** | 437-447, 477-487, 503-510 |
| **$SK150_{BR}$** | 296-306, 336-346, 356-366 |

**Table S2: Validated compound list with their respective percentage control**

| Test compound (1x) conc (nM) | Compound Name | Percentage control |
|---|---|---|
| 10000 | VU0490742 | 70% |
| 10000 | VU0514818 | 68% |
| 10000 | VU0624167 | 65% |
| 10000 | VU0534073 | 64% |
| 10000 | VU0617940 | 64% |
| 10000 | VU0079850 | 63% |
| 10000 | VU0617926 | 41% |
| 10000 | VU0284203 | 35% |
| 10000 | VU0645728 | 23% |

**Table S3: Hydrogen bonding residues to each of the nine validated hits**

| Compounds | Protein residues forming | |
|---|---|---|
| | Backbone HBs | Sidechain HBs |
| 1 | 255 | 285,337,367 |
| 2 | 255,367 | 361,367 |
| 3 | 362,366,367 | 253,337, 339 |
| 4 | 255,367 | 255,337,341,356 |
| 5 | 255, 362 | 285,339,340,356,361,367 |
| 6 | - | 255,285,339,361,365,367 |
| 7 | - | 285,337,339,340,356,365,367 |
| 8 | 362 | 285,337,339,367 |
| 9 | 285 | 339,340,361,365,367 |

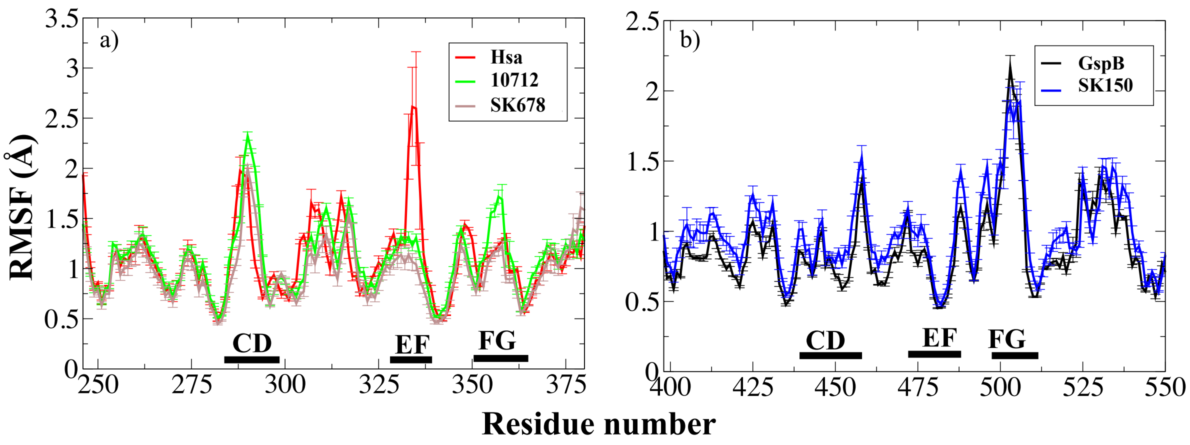


**Fig S1: a) Root mean square fluctuation of $Hsa_{BR}$, $10712_{BR}$ and $SK678_{BR}$ (a); $GspB_{BR}$ and $SK150_{BR}$ (b) from MD simulation showing DC, EF, FG loop regions. The residue numbers of $Hsa_{BR}$ and $GspB_{BR}$ were used in (a) and (b) respectively. The error bar is SEM from three independent simulations.**

a)

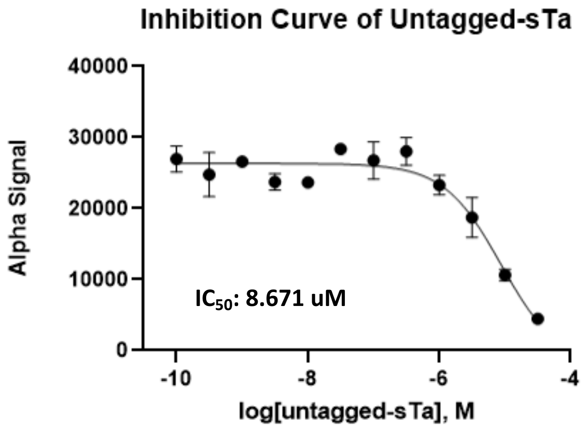


b)

| Log (1x) untagged-sTa (Log M) | (1x) untagged-sTa (nM) | Percentage control |
|---|---|---|
| -4.5 | 30000 | 17% |
| -5.0 | 10000 | 39% |
| -5.5 | 3000 | 69% |
| -6.0 | 1000 | 86% |
| -6.5 | 300 | 104% |
| -7.0 | 100 | 99% |
| -7.5 | 30 | 105% |
| -8.0 | 10 | 88% |
| -8.5 | 3 | 88% |
| -9.0 | 1 | 99% |
| -9.5 | 0.3 | 92% |
| -10.0 | 0 | |

**Fig S2:a) Inhibition curve of untagged sTa (Alpha Signal vs Log (untagged sTa)) used to calculate $IC_{50}$; b) Table showing the percentage control at different concentration of untagged sTa.**

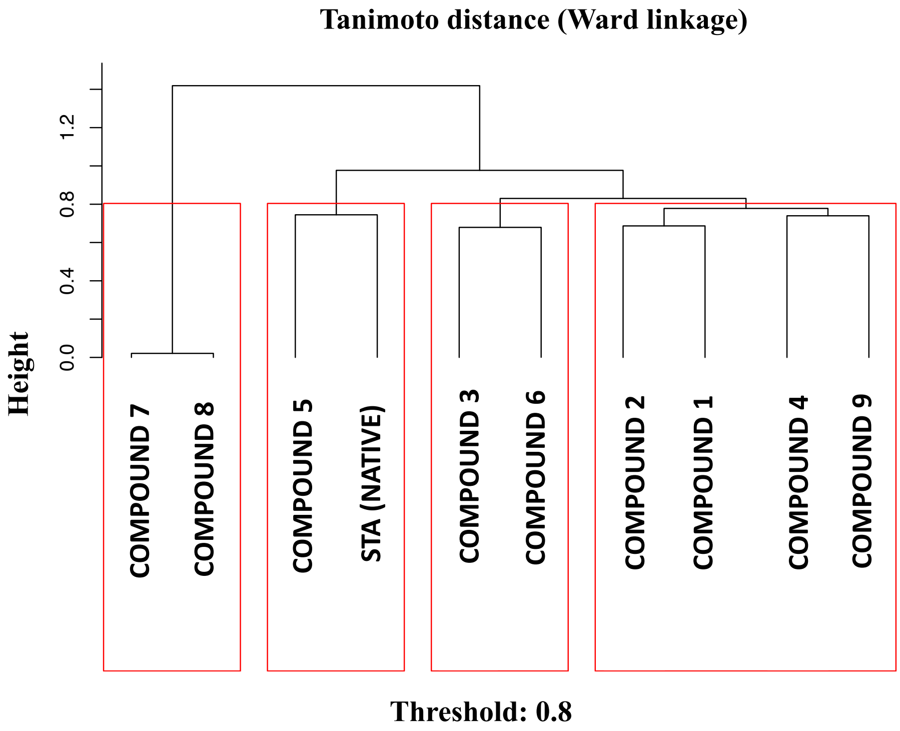


**Fig S3: Hierarchical clustering based on MACCS fingerprint of the nine validated hits and the native ligand**

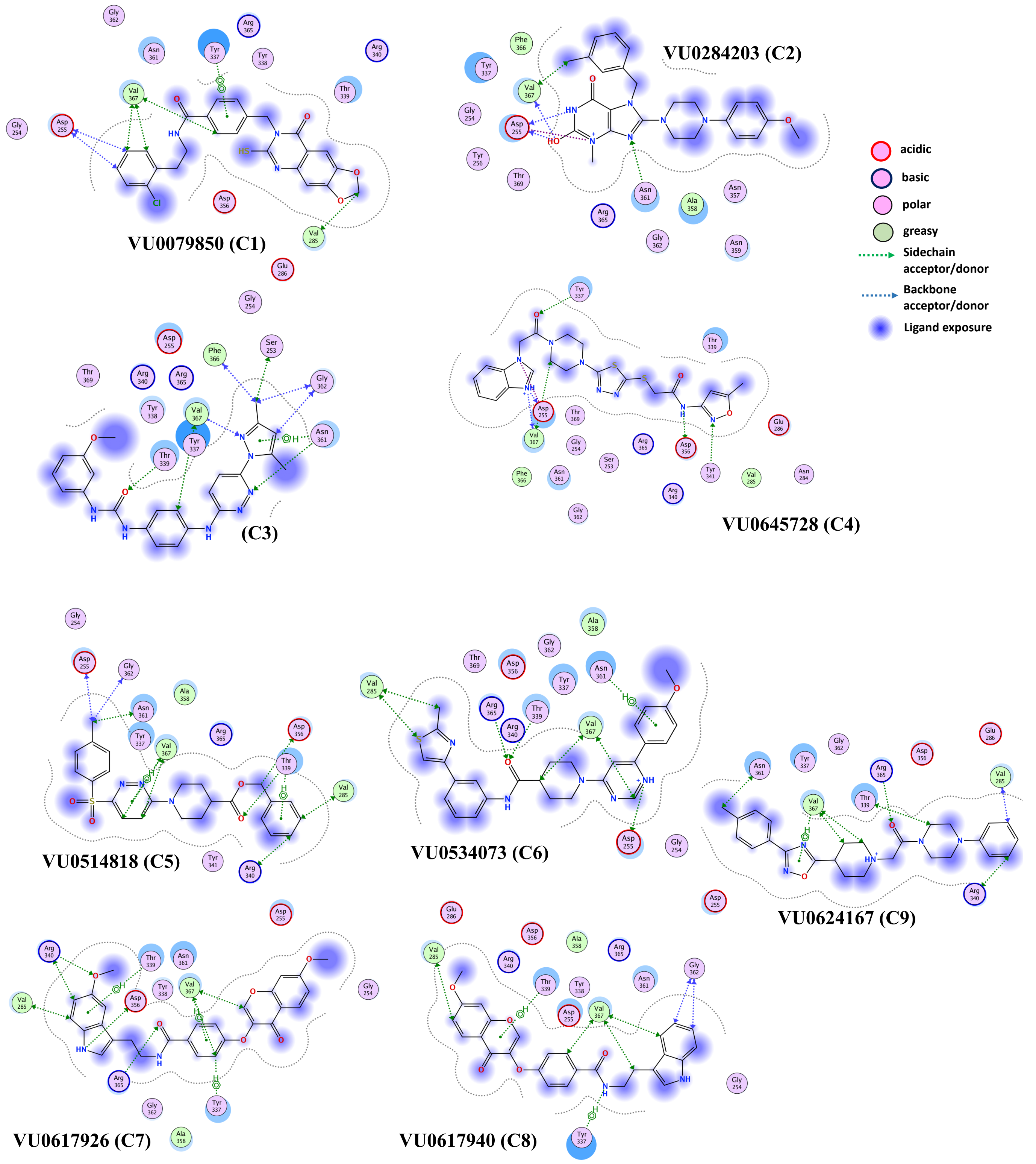


**Fig S4: Interaction map of the nine hits in the binding pocket of the $Hsa_{BR}$**
